# Canonical insulin-like receptor DAF-2 signaling-independent patterning and role for FoxO/DAF- 16 in early embryos

**DOI:** 10.64898/2026.08.04.742880

**Authors:** Michael S. Mauro, Jessica M. Peiser, Sophia L. Martin, J. Tristian Wiles, Julien Dumont, Mimi Shirasu-Hiza, Julie C. Canman

## Abstract

Forkhead box O (FoxO) transcription factors (DAF-16 in *Caenorhabditis elegans*) regulate aging, metabolism, and development. Canonically, FoxO/DAF-16 activity is regulated by insulin/insulin-like receptor (DAF-2) signaling, which inhibits its nuclear localization. In *C. elegans*, strong loss-of-function *daf-16; daf-2* double mutants are embryonic lethal. However, because *daf-2* null mutants are maternally rescued as embryos and then arrest as larvae, the role of DAF-2 signaling in embryogenesis is unknown. We therefore used quantitative imaging and genetics to study DAF- 16 and DAF-2 in early *C. elegans* embryos. DAF-16 was uniformly low in all nuclei at the 2- to 4- cell stage. From the 8- to 64-cell stage, DAF-16 became enriched in 1-2 nuclei of germ lineage cells. This patterning required germ fate determinants and the lipid phosphatase PTEN/DAF-18, but not DAF-2 kinase activity, and was independent of maternal age. We also found that *daf-16; daf-2* double mutant embryos failed morphogenesis, with severe mitotic defects as early as the 1-cell stage. This work identifies germ lineage-specific DAF-16 patterning and a role for DAF-16 in early embryogenesis that is independent of canonical DAF-2 signaling.

## Introduction

Forkhead box O (FoxO) transcription factors are highly conserved from worms to humans and play critical roles in metabolism, cancer, stress tolerance, cell cycle regulation, aging, reproduction, and development.^1–5^ Canonically, FoxO family members are inhibited by the insulin/insulin-like growth factor (IGF-1) signaling (IIS) pathway,^4,6,7^ which was first identified in the model worm *Caenorhabditis elegans.*^8–10^ *C. elegans* has a simplified IIS pathway with a single insulin/IGF-1 receptor (DAF-2) and a single downstream FoxO family transcription factor (DAF- 16).^4^ In contrast, mammals have three insulin/IGF-1 receptor family proteins (INSR, IGF1R, IRR) and four FoxO family members (FoxO1/3/4/6)^2,11^. As in mammals, activation of INSR/DAF-2 inhibits DAF-16 downstream transcriptional activity (**Fig. 1A**).^4,8,10,12–19^ Under nutrient-rich conditions, worm insulin-like peptides^20^ bind to DAF-2, thereby activating its tyrosine kinase activity to stimulate phosphatidylinositol-3 (PI3K; catalytic subunit AGE-1 in worms) kinase activity^21,22^. The PI3K/AGE-1-dependent conversion of PIP_2_ to PIP_3_ activates downstream PDK-1 and AKT-family kinases,^23,24^ leading to phosphorylation of DAF-16, which prevents its nuclear entry and thus its transcriptional activity (**Fig. 1A**). Under starvation conditions, when DAF-2 is inactive, the lipid phosphatase PTEN (DAF-18 in *C. elegans*)^25^ converts PIP_3_ to PIP_2_, reducing PDK-1 and AKT activity and allowing DAF-16 nuclear entry and transcriptional activation^19^ (**Fig. 1A**). Thus, canonically, DAF-2 signaling inhibits DAF-16 nuclear localization and transcription factor activity.

**Figure 1.**
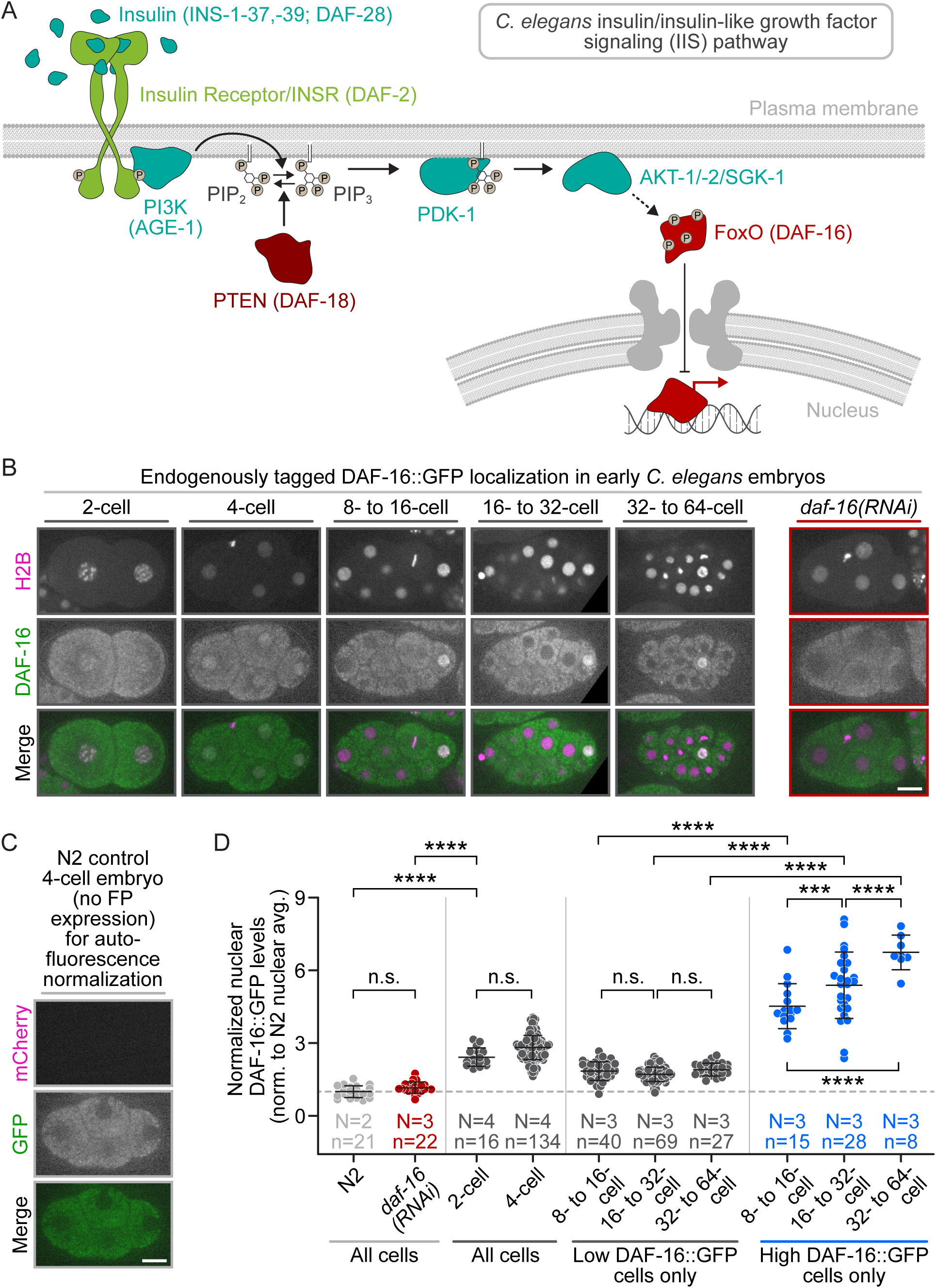
FoxO/DAF-16 patterning in the 8-cell *C. elegans* embryo. **A)** Schematic depicting the worm IIS pathway; worm protein names in parentheses. **B)** Endogenously tagged DAF- 16::GFP (green) and mCherry::H2B (magenta) localization in control 2- through 64-cell embryos and *daf-16(RNAi)* 4-cell embryos. **C)** Autofluorescence levels in a N2 4-cell embryo with same imaging/display settings as in **B**. **(B, C)** Black box behind some images for display; scale bars=10 μm. **D)** Graph plotting average nuclear DAF-16::GFP levels in *daf-16(RNAi)* and control embryos normalized to average N2 4-cell nuclear autofluorescence levels. Error bars=SD; N/n=number of experimental replicates/nuclei scored per genotype, developmental stage, or relative DAF- 16::GFP levels by color (gray=low DAF-16::GFP levels, red=*daf-16(RNAi)*, blue=high DAF- 16::GFP levels) ; n.s.=p-value not significant (>0.05), ***=p-value ≤0.001, ****=p-value ≤0.0001 (1-way ANOVA; only relevant comparisons shown; see **Table S1**).

In canonical IIS, because DAF-2 inhibits DAF-16, genetic disruption of *daf-2* typically has no phenotypic effect if *daf-16* is co-disrupted. For example, severe loss-of-function *daf-2* mutations lead to a developmental arrest due to constitutive dauer larvae formation (Daf-c), and hypomorphic mutations increase adult lifespan.^9,26–28^ Both phenotypes are suppressed (or lost) in *daf-2*; *daf-16* double mutants, suggesting that that DAF-16 is regulated by canonical IIS signaling in these contexts.^9,21,26–30^ Similarly, DAF-2 signaling functions through DAF-16 to promote germline proliferation and oocyte quality in adults^31–33^ and to promote the starvation-induced cell cycle arrest of somatic cells in larvae^34^. In these canonical IIS regulated-processes, genetic co-disruption of *daf-16* blocks the phenotypic effects of *daf-2* disruption.

On the other hand, DAF-2 and DAF-16 can function independently or redundantly wherein *daf-2* loss-of-function mutant phenotypes are not blocked by *daf-16* co-disruption. For example, DAF-2 signaling does not act through DAF-16 to stall meiotic progression and oocyte production in adults under starvation conditions or to promote the starvation-induced cell cycle arrest of larval germline precursor cells.^35–37^ In these contexts, the phenotypes caused by disruption of *daf-*2 are not affected by co-disruption of *daf-16*.^35–37^ DAF-2 and DAF-16 can also function independently or redundantly in the same biological process, most dramatically during embryogenesis. DAF-2 and DAF-16 are both expressed in early embryos^38,39^ and reducing DAF-2 function leads to some embryo and larval lethality^26–28,40^, but due to maternal rescue of *daf-2* null embryos followed by larval arrest, the role of DAF-2 in embryogenesis is not known. Nonetheless, strong loss-of-function *daf-16; daf-2* double mutants produce only dead embryos at 25°C.^26–28^ As *daf-16* null mutants alone produce viable embryos,^27,29^ this synthetic lethality suggests that DAF-16 has an embryonic role that is distinct from canonical DAF*-2* signaling.

Although the role of DAF-16 has been extensively studied in larval and adult stages, its regulation and function during embryogenesis is not understood. Here, we investigated DAF-16 in early *C. elegans* embryogenesis using a combination of live-cell imaging and lineage tracing, quantitative image analysis, and genetic approaches. We examined the patterning of DAF-16 nuclear localization in embryos during early cleavage stage development and assessed its regulation by IIS pathway components (**Fig. 1A**). Our findings reveal germ lineage-specific patterning of DAF-16 starting at the 8-cell stage and uncover a critical role for DAF-16 in early embryogenesis that acts in parallel to or redundantly with canonical DAF-2 signaling.

## Results

### DAF-16 exhibits a specific pattern of nuclear enrichment in germ precursor cells during early embryogenesis

Active FoxO/DAF-16 is known to localize to the nucleus (**Fig. 1A**). Because the pattern of expression and localization of DAF-16 during early *C. elegans* development had not been characterized, we first imaged DAF-16::GFP in oocytes and early cleavage stage embryos using spinning disk confocal microscopy at 25°C. For this, we generated a strain co-expressing endogenously tagged DAF-16::GFP^41^ and mCherry::Histone2B^42^ (mCh::H2B) to label nuclei. Because the DAF-16::GFP signal was relatively dim, we controlled for background autofluorescence in the GFP channel by performing parallel analyses in wild type (N2 Bristol) embryos and normalizing fluorescence intensity across samples. Consistent with prior reports^39^, DAF-16 was enriched in nuclei of immature oocytes (−5+ oocytes) but was not detectable following oocyte maturation (−1 to −4 oocytes) (**Fig. S1**). No nuclear DAF-16 was detected in germline cells, oocytes, or embryos after *daf-16(RNAi)*, confirming specificity (**Fig. 1B-D, Fig. S1**). In early embryos at the 2- to 4-cell stage, DAF-16::GFP showed low but uniform nuclear enrichment in all cells (**Fig. 1B-D**, **S2**). In contrast, in 8-cell stage embryos, we no longer observed nuclear DAF-16 in anterior embryonic cells but observed increased nuclear levels in 1 or 2 posterior embryonic cells (**Fig. 1B-D**). From the 16- to 64-cell stages, this DAF-16 nuclear enrichment remained restricted to 1 or 2 posterior cells and increased as cell and nuclear volume decreased with each cleavage division^43–45^ (**Fig. 1B-D**). These results suggest that, at the 8-cell stage, nuclear DAF-16 is preferentially enriched in a posterior cell lineage during early worm embryogenesis.

The *C. elegans* embryonic cell lineage map (**Fig. 2A**) is well characterized and highly invariant, allowing individual cells to be readily identified by time-lapse microscopy.^46^ To identify the posterior cell lineage with elevated nuclear DAF-16, we performed time-lapse cell lineage tracing. We used mCh::H2B to monitor each cell nucleus from the 4-cell to 32-cell stage (**Video S1**) and then imaged endogenously tagged DAF-16::GFP after each round of cleavage division. We found that nuclear DAF-16 levels were ∼1.5 fold higher in the P3 and C cells (8- to 16-cell stage) and ∼3 fold higher in P4 and D cells (16- to 32-cell stage) compared with anterior somatic cells (ABal/r (ABax; 8- to 16-cell) and ABal/ra/p (ABaxx; 16- to 32-cell), respectively) (**Fig. 2B-E, Video S1**). These results are consistent with our earlier observation that nuclear DAF-16 enrichment increases during successive cleavage divisions as nuclear volume decreases with cell size^43–45^ (**Fig. 1B-D**).

**Figure 2.**
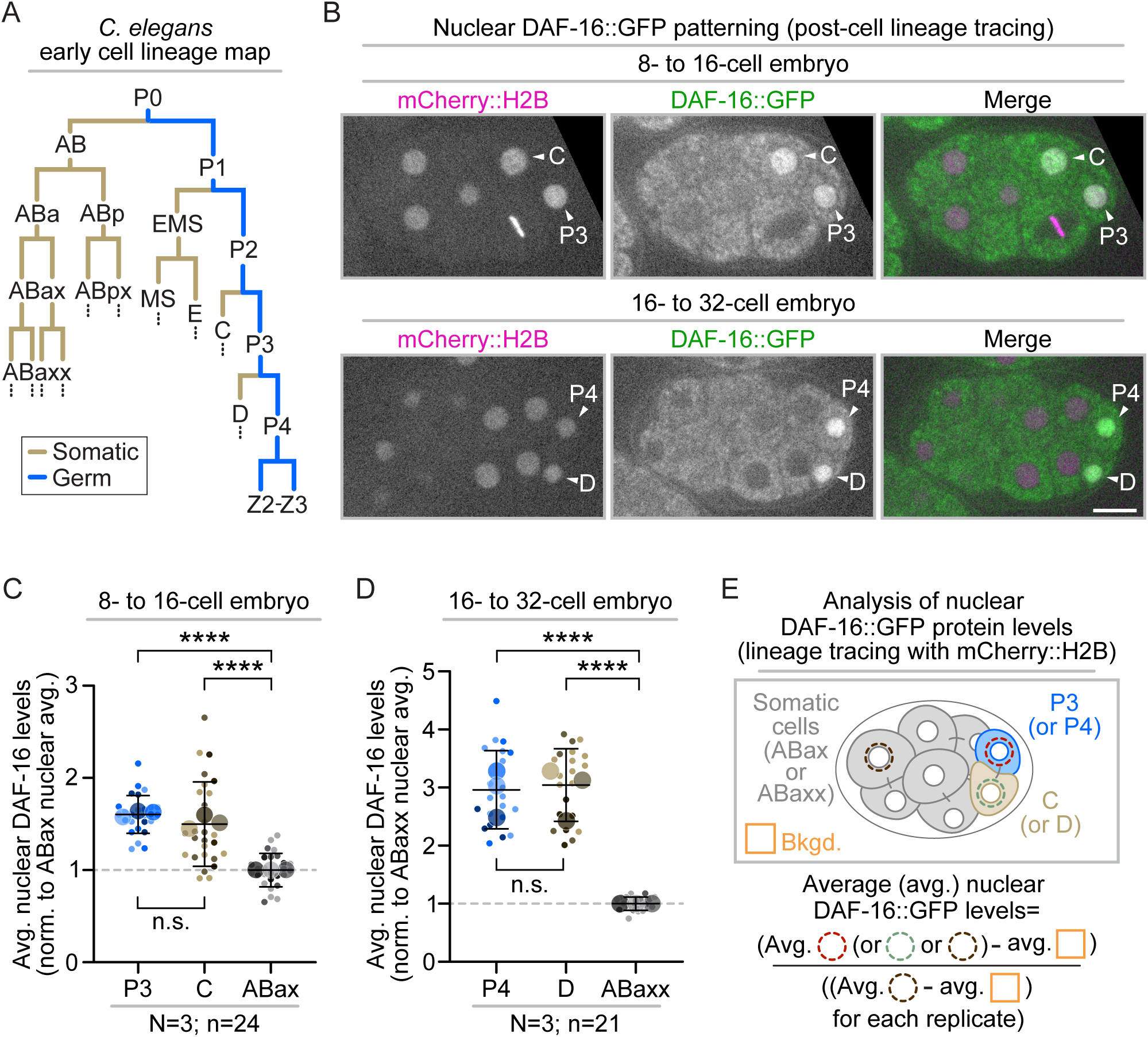
Nuclear FoxO/DAF-16 is enriched specifically in germ lineage cells. **A)** Schematic depicting *C. elegans* early embryonic somatic (tan) and germ (blue) cell lineage map. **B)** Representative images of DAF-16::GFP (green) and mCherry::H2B (magenta) localization in the P3 and C cells (arrowheads, top panels) and P4 and D cells (arrowheads, bottom panels) after lineage tracking with mCherry::H2B (**Video S1**); black box behind some images for display; scale bar=10 μm. **C-D)** Graphs plotting the average nuclear levels of DAF-16::GFP in **(C)** 8- to 16-cell embryos and **(D)** 16- to 32-cell embryos; results normalized to average nuclear levels in **(C)** ABax or **(D)** ABaxx; small circles=average levels in individual nuclei, large circles=replicate averages, shading=results from same replicate (*e.g.*, darkest shades from same replicate). Error bars=SD; N/n=number of experimental replicates/number of nuclei scored per cell type; n.s.=p-value not significant (>0.05), ****=p-value ≤0.0001 (1-way ANOVA; see **Table S1**). **E)** Schematic depicting analysis in **C-D**.

The posterior cells with nuclear DAF-16 enrichment are pluripotent and divide asymmetrically to form distinct cell lineages. P3 and P4 cells form the germ lineage (P lineage), whereas their larger sister cells, C and D, form somatic lineages (**Fig. 2A**). Notably, DAF-16 was not observed in the daughters of C or D cells (**Fig. 2B**), suggesting that it is subsequently degraded or excluded from nuclei following cell division. Together, these findings indicate that nuclear-enriched DAF-16 is preferentially inherited by the germ lineage, which ultimately gives rise to all gametes in the adult worm^46^.

### Germ fate determinants regulate nuclear DAF-16 in the P3 germ precursor cell

Having identified the germ precursor cells as the embryonic lineage exhibiting nuclear DAF-16 enrichment, we next tested if germ cell fate is required for this patterning. In *C. elegans*, germ cell fate is specified by the CCCH (three cysteines and a histidine)-type zinc-finger protein, PIE-1 (Pharynx and Intestine in Excess)^47–51^, along with the related factor POS-1 (Posterior Segregation)^52^, which is required for normal PIE-1 localization^53,54^ (**Fig. 3A**). Both PIE-1 and POS- 1 are asymmetrically inherited by germ precursor cells (P lineage) and regulate key germ cell-specific processes, including transcription and translational control.^47–50,53–64^

**Figure 3.**
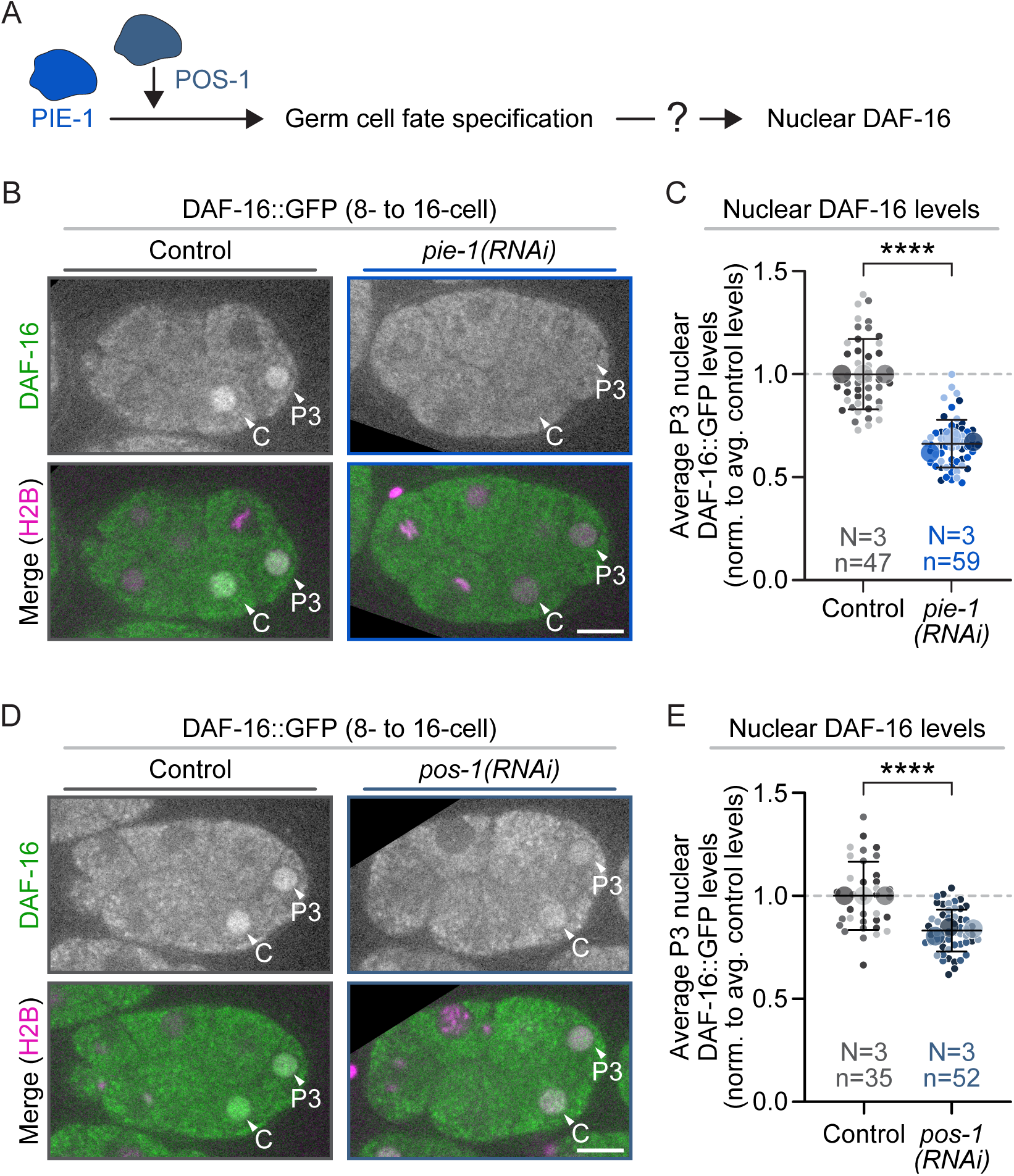
Nuclear DAF-16 germ precursor cell enrichment is germ fate-dependent. **A)** Schematic depicting the role of PIE-1 and POS-1 in germ cell fate specification and potential upstream role of DAF-16::GFP. **B, D)** Representative images of DAF-16::GFP (green) and mCherry::H2B (magenta) localization in embryos with and without **(B)** *pie-1(RNAi)* or **(D)** *pos- 1(RNAi).* Black box behind some images for display; arrowheads indicate P3 and C cells; scale bars=10 μm. **C, E)** Graphs plotting average P3 nuclear levels of DAF-16::GFP in control (grays) embryos with and without **(C)** *pie-1(RNAi)* (royal blues), and **(E)** *pos-1(RNAi)* (dusty blues). **C, E)** Results normalized to average nuclear levels in controls; small circles=average levels in individual nuclei, large circles=replicate averages, shading=results from same replicate (*e.g.*, darkest shades for each genotype from same replicate). Error bars=SD; N/n=number of experimental replicates/number of nuclei scored per genotype by color; ****=p-value ≤0.0001 (Student’s t-test, unpaired; see **Table S1**).

To test whether these determinants are required for DAF-16 patterning, we measured nuclear DAF-16::GFP levels in the P3 cell following RNAi-mediated depletion of PIE-1 or POS-1, which greatly reduces protein levels and leads to embryonic lethality.^65^ Compared with control embryos, nuclear DAF-16 levels were reduced by ∼30-38% in *pie-1(RNAi)* embryos and ∼15-20% in *pos-1(RNAi)* embryos **(Fig. 3B-E**). We next tested if DAF-16 reciprocally regulates PIE-1. Although DAF-16 is not essential for germ fate specification,^33^ it is reported to regulate PIE-1 expression during aging^66^ (however, for contradictory results see^67^). Thus, we quantified the levels of endogenously tagged PIE-1::GFP^68^ in germ precursor cells (P2, P3, and P4 (4-cell to ∼32-cell stages)) with and without *daf-16(RNAi)*. We did not observe any significant difference in PIE-1 levels or localization between control and *daf-16(RNAi)* embryos at any stage (**Fig. S3**). These results suggest that proper germ cell fate specification is required for nuclear enrichment of DAF- 16 (or its protein stability) in the germ lineage but not *vice versa*.

### P3 nuclear DAF-16 requires DAF-18 and AGE-1 but is independent of DAF-2

FoxO/DAF-16 is regulated by several different pathways.^4,6,69^ We next tested if DAF-16 patterning during embryogenesis is regulated by the canonical IIS pathway (**Fig. 1A**) using both RNAi depletion and classic genetic alleles. As above, all imaging was done at 25°C because many DAF-2 dependent processes are temperature-sensitive (*e.g.*, see^27,29,70^). We first depleted different IIS pathway components using dsRNA injection-based RNAi-mediated depletion and quantified the average nuclear levels of DAF-16::GFP in the P3 cell (8- to 16-cell stage), with and without RNAi knockdown of each IIS gene (**Fig. 4A-B**). RNAi knockdown was confirmed by analysis of nuclear DAF-16 levels in intestinal cells, which is blocked by IIS activity and thus increases upon disruption of IIS^19,27,71,72^ (**Fig. S4A, C**), and by quantitative analysis of specific endogenously tagged fluorescent reporters, when available (^73^ and Glow Worm project; **Fig. S5A- B**). RNAi knockdown of *daf-2* did not affect P3 nuclear DAF-16 levels (**Fig. 4A-B**), suggesting that this early DAF-16 patterning is DAF-2 activity-independent. In contrast, RNAi knockdown of PI3K/AGE-1 significantly reduced P3 nuclear DAF-16 levels (20-25% lower than in controls, **Fig. 4A-B**). RNAi knockdown of PTEN/DAF-18 also decreased P3 nuclear DAF-16 levels (16-25% lower than in controls, **Fig. 4A-B**). Together, these results suggest that DAF-16 germ lineage patterning is controlled by phospholipid regulators in the IIS pathway but is potentially independent of insulin-like receptor signaling upstream of these phospholipid regulators.

**Figure 4.**
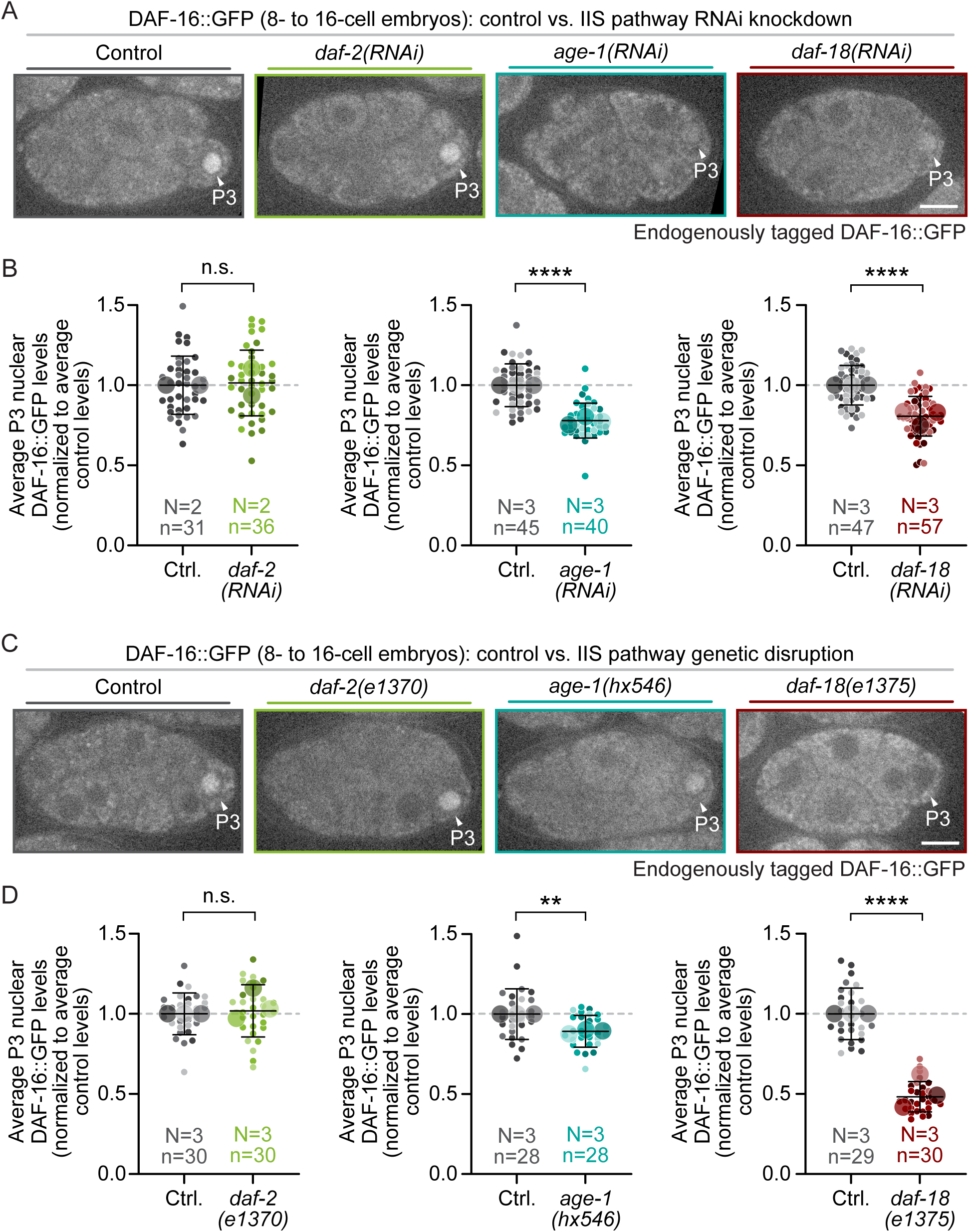
P3 FoxO/DAF-16 nuclear enrichment requires PTEN/DAF-18 and to a lesser extent PI3K/AGE-1 but is INSR/DAF-2 independent. **A, C)** Representative images of P3 nuclear DAF- 16::GFP localization with and without **A**) RNAi-knockdown or **C**) genetic disruption of indicated IIS pathway genes (see **Fig. 1A**); arrowheads indicate P3 cells; black box behind some images for display; scale bars=10 μm. **B**) Graphs plotting average P3 nuclear levels of DAF-16::GFP in embryos from control (grays), *daf-2(RNAi)* (greens), *age-1(RNAi)* (teals), and *daf-18(RNAi)* (reds) worms. **D**) Graphs plotting average P3 nuclear levels of DAF-16::GFP in embryos from control (grays), *daf-2(e1370)* (greens), *age-1(hx546)* (teals), or *daf-18(e1375)* (reds) worms. (**B, D**) Results normalized to average nuclear levels in controls; small circles=average levels in individual nuclei, large circles=replicate averages, shading=results from same replicate (*e.g.*, darkest shades from same replicate). Error bars=SD; N/n=number of experimental replicates/number of nuclei scored per genotype by color; n.s.=p-value not significant (>0.05), **=p-value ≤0.01, ****=p- value ≤0.0001 (Student’s t-test, unpaired; see **Table S1**).

To confirm our RNAi results on IIS genes in FoxO/DAF-16 patterning, we examined the effect of classic IIS pathway loss-of-function genetic alleles using strains containing these alleles and expressing endogenously tagged DAF-16::GFP. We again confirmed IIS pathway disruption by quantitative analysis of nuclear DAF-16 levels in intestinal cells (**Fig. S4B-C**). To disrupt INSR/DAF-2 function, we used the *daf-2(e1370)* allele, which contains a missense mutation in the kinase domain that blocks kinase activity^74^ and doubles worm lifespan,^9^ but does not lead to significant embryonic lethality.^26,75^ Similar to our results with *daf-2(RNAi)* (**Fig. 4A-B**), P3 nuclear DAF-16 levels were not affected in *daf-2(e1370)* embryos (**Fig. 4**). To disrupt PI3K/AGE-1 activity, we used the *age-1(hx546)* allele,^76^ which contains a missense mutation in the kinase domain that increases lifespan by ∼65%.^22^ Again, consistent with RNAi of *age-1* (**Fig. 4A-B**), P3 nuclear DAF- 16 levels were reduced by the *age-1(hx546)* allele, although to a lesser extent (10-13% lower than in controls, **Fig. 4C-D**). To disrupt PTEN/DAF-18 activity, we used the *daf-18(e1375)* allele, which contains a small insertion with a premature stop codon that disrupts the carboxyl-terminal half of the protein and reduces the Daf-c and lifespan extending effects of AGE-1 or DAF-2 inhibition.^21,25,28,30,77–79^ Similar to our results with *daf-18(RNAi)* (**Fig. 4A-B**), P3 nuclear DAF-16 levels were reduced in *daf-18(e1375)* embryos to an even greater extent (38-58% lower than in controls, **Fig. 4C-D**). Taken together, our genetic analysis of the IIS pathway is consistent with our RNAi analysis and suggests that phospholipid regulation, rather than canonical insulin-like receptor kinase signaling, is the primary driver of nuclear DAF-16 patterning in the germ lineage.

### DAF-16 embryonic patterning is maternal age-independent

Like in female humans, worms undergo maternal reproductive cessation midlife (day ∼7 of adulthood)^80,81^ and maternal reproductive aging is regulated by DAF-2 and DAF-16 activity.^82^ Somatic aging is reported to increase nuclear DAF-16 levels in the intestine,^83^ so we reasoned that maternal age may impact DAF-16 levels in early embryos. To test the role of maternal age in DAF-16 embryonic patterning, we first mated worms to ensure sperm number was not limiting^82^ and then quantified endogenously tagged DAF-16::GFP levels in P3 nuclei in embryos from young (1-day old adult) vs. aged (6-day old adult) mated worms (**Fig. 5A**). We found no difference in P3 DAF-16 nuclear levels between embryos from young and aged mothers (**Fig. 5B-C**). Thus, P3 nuclear DAF-16 levels in the early embryo are not significantly impacted by maternal reproductive age.

**Figure 5.**
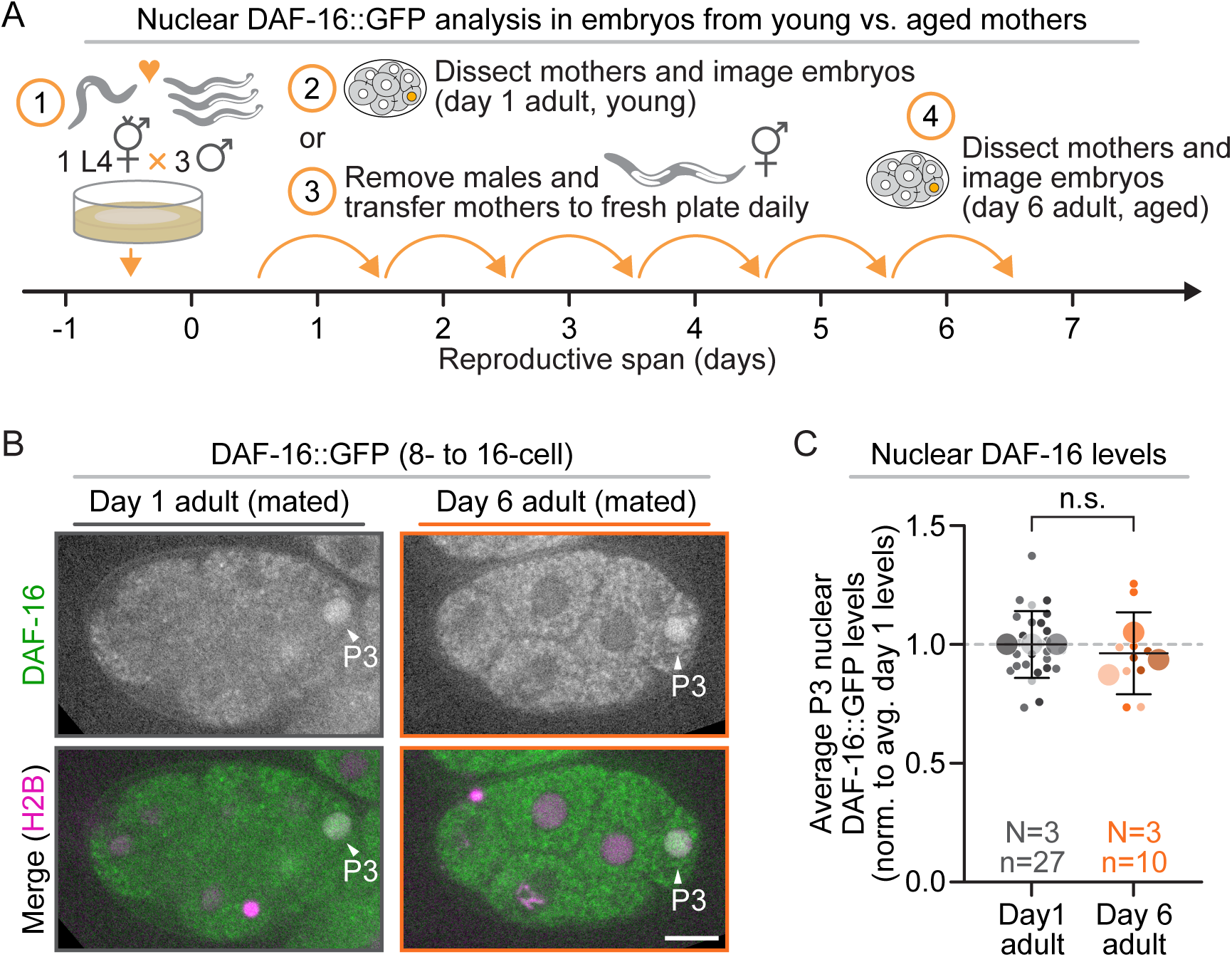
P3 FoxO/DAF-16 nuclear enrichment is maternal age-independent. **A)** Schematic of experimental design. L4 worms were singled and either mated for 1 day then dissected for embryo imaging or transferred each day for a total of 6 days prior to embryo imaging. **B)** Representative images of P3 nuclear DAF-16::GFP localization from young (day 1 of adulthood) vs. aged (day 6 of adulthood) mothers; arrowheads indicate P3 cells; scale bar=10 μm. **C)** Graph plotting average P3 nuclear levels of DAF-16::GFP in embryos from young (grays) and aged (oranges) mothers. Results normalized to average nuclear levels in embryos from young mothers; small circles=average levels in individual nuclei, large circles=replicate averages, shading=results from same replicate (*e.g.*, darkest shades from same replicate). Error bars=SD; N/n=number of experimental replicates/number of nuclei scored per maternal age by color; n.s.=p-value not significant (>0.05) (Student’s t-test, unpaired; see **Table S1**).

### FoxO/DAF-16 and INSR/DAF-2 co-disruption leads to defects in morphogenesis and early mitoses

It has long been known that genetic co-disruption of *daf-16* and *daf-2* activity using severe loss-of-function alleles leads to 100% embryonic lethality at 25°C (vs. ∼39% at 15°C^27^),^26–28^ but the exact timing and cause of this lethality remains unknown. To confirm this synthetic embryonic lethality, we first performed brood size and embryonic lethality assays in *daf-16(mgDf50)*; *daf- 2(m65)* double mutant worms. *daf-16(mgDf50)* is a null allele^84^ and *daf-2(m65)* is a nonsense allele with a premature stop codon that removes part of the kinase domain and the carboxyl-terminal intracellular region.^26,27,85^ We were unable to test the *daf-2(m65)* allele alone without *daf- 16* co-disruption. The *daf-2(m65)* allele must be maintained over a balancer and maternally provided *daf-2* rescues *daf-2(m65)* homozygous embryos; the embryos hatch but enter a Daf-c larval arrest that blocks development before reproduction.^26^ As expected, we found that *daf- 16(mgDf50); daf-2(m65)* double mutants had a significantly lower total and daily brood size than either *daf-16(mgDf50)* single mutants or controls (**Fig. 6A**). *daf-16(mgDf50); daf-2(m65)* double mutants produced mostly dead embryos and larva after either 24 hours or 4 hours at 25°C, whereas *daf-16(mgDf50)* single mutants produced mostly viable progeny^27^ (**Fig. 6B**). Thus, our results suggest, as was previously reported,^26–28^ severe co-disruption of *daf-16* and *daf-2* leads to a significant reduction in brood size and high embryonic lethality.

**Figure 6.**
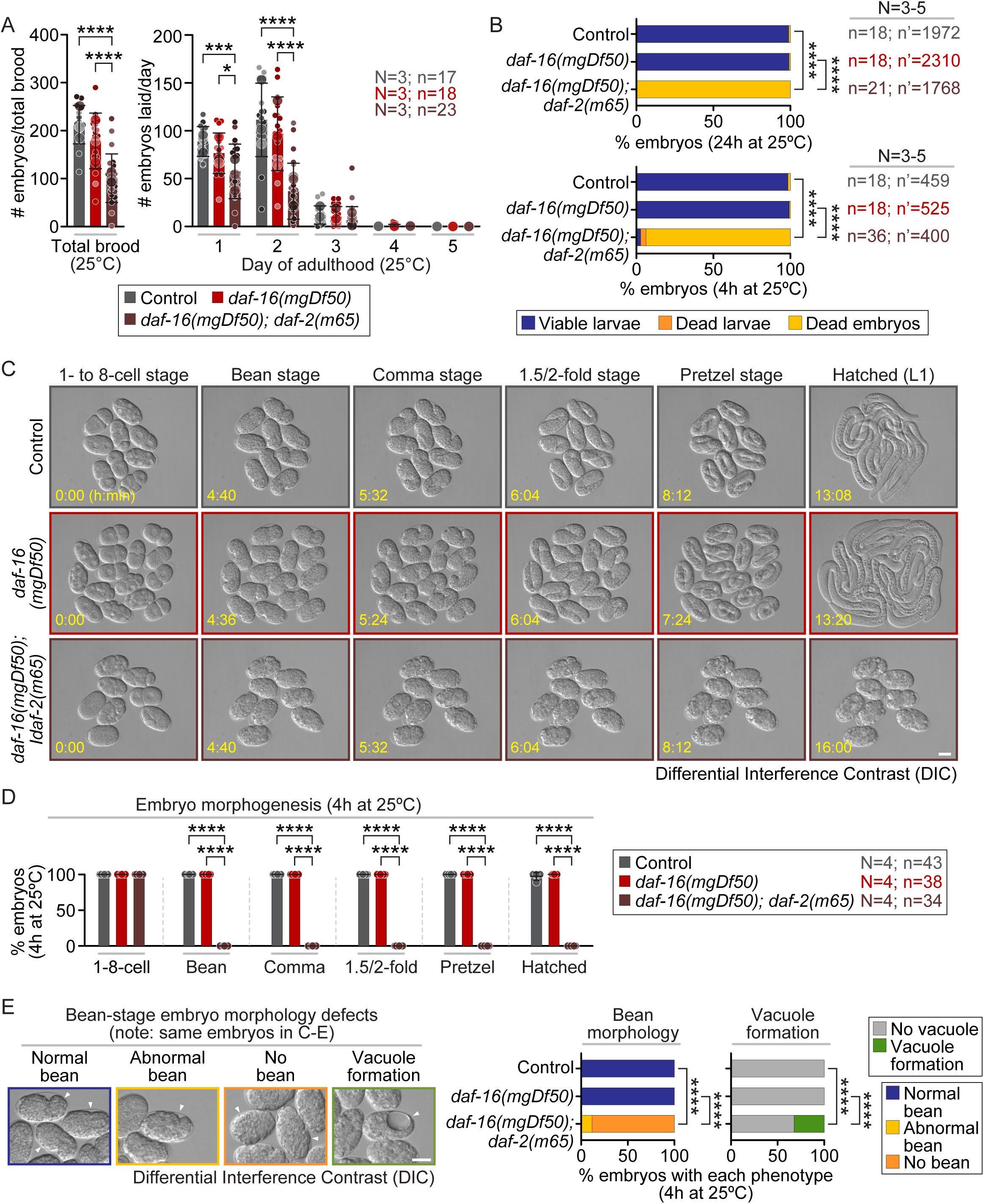
Embryos from severe *daf-16; daf-2* double mutants fail to undergo morphogenesis. **A)** Total brood size (left) and number of embryos laid per day 1-5 of adulthood (right) at 25°C for unmated control (grey), *daf-16(mgDf50)* (red), and *daf-16(mgDf50); daf-2(m65)* (brown) double mutant worms. N/n=number of experimental replicates/number of worms scored per genotype by color. **B)** Percent of embryos that develop into viable larvae (blue), undergo larval lethality (orange), or die as embryos (yellow) after 24 hours (top) and 4 hours (bottom) at 25°C from control, *daf-16(mgDf50)*, and *daf-16(mgDf50); daf-2(m65)* double mutant worms. N/n/n’=number of experimental replicates/number of worms/number of embryos scored per genotype by color. **C)** Representative images from Differential Interference Contrast (DIC) time-lapse image series of early embryo development in control (top row), *daf-16(mgDf50)* (center row), and *daf-16(mgDf50); daf-2(m65)* double mutants (bottom row) after 4 hours at 25°C; time in hours:minutes indicated in yellow; scale bar=20 μm. **D)** Percent of embryos that develop to the 1- to 8-cell, bean, comma, 1.5/2-fold, pretzel stage, and hatching for control (grey), *daf-16(mgDf50)* (red), and *daf-16(mgDf50); daf-2(m65)* double mutants (brown); N/n=number of experimental replicates/number of embryos scored per genotype by color. **E)** Examples of normal bean formation (blue), abnormal bean formation (yellow), no bean formation (orange), and vacuole formation (green) in embryos from control, *daf-16(mgDf50)*, and *daf-16(mgDf50); daf-2(m65)* double mutants; white arrowheads=examples of embryos with each phenotype; scale bar=20 μm. Note: same dataset analyzed in **C-E**. (**A-B, D-E**) *=p-value ≤0.05, ***=p-value ≤0.001, ****=p- value ≤0.0001 **(A)** (left) 1-way ANOVA, (right) mixed effects ANOVA, **(B, D)** Fisher’s exact test,

We next sought to determine the terminal phenotype for these double mutant embryos. We used time-lapse Differential Interference Contrast (DIC, transmitted light) microscopy with 4-minute resolution after 4 hours at 25°C to image embryo development from the ∼1- to 8-cell stage through hatching (or for ∼16 hours upon failure to develop) and then quantified any observed developmental defects. As expected from the low levels of embryonic lethality, most control and *daf-16(mgDf50)* single mutant embryos progressed normally through the bean, comma, 1.5- to 2-fold, and pretzel (3-fold) stages of development and successfully hatched into larvae (**Fig. 6C-E**, **Video S2**). In contrast, most *daf*-*16(mgDf50); daf*-*2(m65)* double mutant embryos failed to undergo morphogenesis and did not hatch, although they sometimes formed abnormal bean-like morphologies (**Fig. 6C-E**, **Video S2**). About a third of *daf*-*16(mgDf50); daf*-*2(m65)* embryos also formed abnormal blastocoel- or vacuole-like structures within the embryonic cell mass (**Fig. 6E**). Importantly, this terminal phenotype occurs much earlier in development than in single *daf*-*2(e979, m579, or m41)* mutants, which have ∼3-10% embryonic lethality due to late developmental defects (during elongation).^40^ Thus, severe co-disruption of *daf*-*16 and daf*-*2* blocks normal embryo development and morphogenesis.

Finally, to characterize the cause of these developmental defects, we performed 4-D time lapse imaging with 30-second time resolution of the first three mitotic cell divisions in embryos expressing mCh::H2B^86^ to label the chromatin. Again, due to the Daf-c larval arrest phenotype,^26^ the *daf*-*2*(*m65*) allele could not be tested without *daf-16* co-disruption. While control and *daf*-*16*(*mgDf50)* single mutant embryos had little to no observable mitotic defects, *daf*-*16(mgDf50); daf*-*2(m65)* double mutant embryos had major mitotic defects as early as the 1-cell stage, including abnormal metaphase plate formation with mis-aligned chromosomes, anaphase chromosomal bridge formation, and multinucleation (**Fig. 7A-B**, **Video S3**). These results suggest that DAF-2 and DAF-16 play redundant or parallel roles required to maintain genome stability and normal development during early embryogenesis in *C. elegans* (**Fig. 7C**).

**Figure 7.**
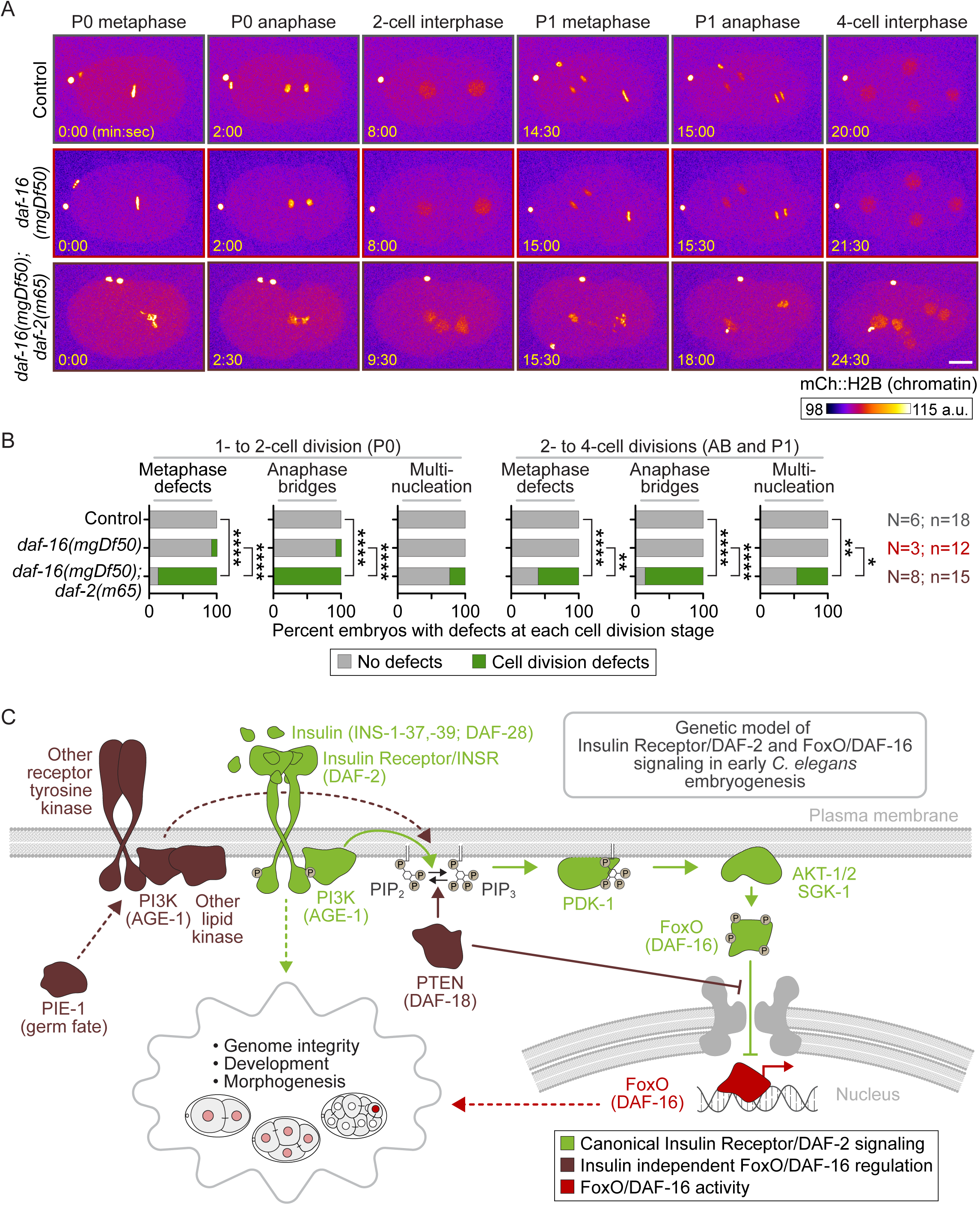
Embryos from severe *daf-16; daf-2* double mutants have genomic instability. **A)** Representative images from time lapse image series of mCherry::histone2B (mCh::H2B) expressing control (top row), *daf-16(mgDf50)* (center row), and *daf-16(mgDf50); daf-2(m65)* double mutant (bottom row) early embryos undergoing the first few cleavage divisions after 4 hours at 25°C; time in minutes:seconds indicated in yellow; scale bar=10 μm. **B)** Percent of embryos from **(A)** with no cell division defects (gray) or defects (green) in metaphase plate formation, anaphase chromosome segregation, and multinucleation in the 1- to 2-cell (P0 division, left) and 2- to 4-cell (AB and P1 divisions, right) divisions; *=p-value ≤0.05, **=p-value ≤0.01, ****=p-value ≤0.0001 (Fisher’s exact test; only statistically significant comparisons shown; see **Table S1**). **C)** Schematic depicting a genetic model for the role for and regulation of DAF-2 and DAF-16 in early embryonic development showing canonical DAF-2 signaling (green), DAF-16 activity (red), and canonical insulin signaling independent or redundant DAF-16 regulation (brown); dashed lines indicate hypothetical regulators and functions.

## Discussion

Taken together, our results show that FoxO/DAF-16 is expressed in early *C. elegans* embryos and functions redundantly with or in parallel to INSR/DAF-2 signaling. While initially uniform in all cell nuclei in very early embryos, starting at the 8-cell stage, we found that DAF-16 was specifically enriched in germ lineage cell nuclei. This germ cell patterning of DAF-16 required the germ cell fate determinants PIE-1 and POS-1, as well as the insulin-signaling components PTEN/DAF-18 and to a lesser extent PI3K/AGE-1, but not DAF-2 kinase activity. Furthermore, while DAF-16 disruption alone did not lead to embryonic lethality, co-disruption of *daf*-*16* and *daf*-*2* using strong loss-of-function alleles blocked morphogenesis and led to severe mitotic defects in the first few cleavage divisions. Together, these results suggest that DAF-16 and DAF-2 function in parallel or redundantly to ensure cell division accuracy and protect genomic integrity in early embryos (**Fig. 7C**).

We were initially surprised that RNAi knockdown of either the phospholipid kinase PI3K/AGE-1 or the phospholipid phosphatase PTEN/DAF-18, which are mechanistically opposed to each other in canonical IIS (*e.g.*, see **Fig. 1A**), led to a similar reduction in P3 nuclear DAF-16 levels. But our analysis of *age-1(hx546)* embryos, which have a point mutation that specifically impacts kinase activity, indicated a minor role for kinase activity and is consistent with a model in which AGE-1 acts more like a scaffold or platform to bring proteins together in early embryonic DAF-16 patterning, as has been found in other contexts.^87–92^ Alternatively, DAF-18 could function as a protein phosphatase rather than a lipid phosphatase.^93,94^ Interestingly, like severe loss-of-function *daf-16; daf-2* double mutants, *daf*-*18; daf*-*2* double mutants also have a high rate of embryonic lethality (70%).^27^ Thus, we predict that *daf-18; daf-2* embryos will also have defects in genetic stability. It remains unclear what functions upstream of DAF-18 to control DAF-16 patterning in the early germ precursor cells. In other contexts, DAF-16 is regulated by JNK (c-Jun N-terminal kinase), AMPK (AMP-activated protein kinase), TOR (target of rapamycin), Ras/ERK (Extracellular signal-regulated kinase), CAMKII (Ca^2+^/calmodulin-dependent kinase type II), and nuclear hormone receptor (DAF-12) signaling pathways.^16,17,35,69,95,96^ Identifying the upstream regulator(s) of PTEN/DAF-18-dependent DAF-16 patterning will be an important goal for future studies.

What drives FoxO/DAF-16 inheritance and maintenance in germ lineage cells? The early worm embryo undergoes asymmetric cell divisions starting at the 1-cell stage. This asymmetric cell division is driven by the conserved cell polarity machinery, which polarizes the 1-cell embryo (zygote) along the anterior-posterior axis to produce asymmetric daughter cells with different cell fate determinants, such as PIE-1 (for review see^97,98^). While we do not know why DAF-16 is uniformly distributed in 2- to 4-cell embryos, PIE-1 is also dispensable for *nanos-related/nos-2* mRNA localization in 4-cell stage embryos but is required for *nos-2* mRNA enrichment in the P3 germ precursor cell at the 8-cell stage,^54^ suggesting this delay in PIE-1-dependent patterning is not specific to DAF-16. It seems likely that protein degradation also controls DAF-16 patterning; indeed, DAF-16 degradation is important in *C. elegans* lifespan regulation^99–101^ and human FoxO3 is degraded in a cell cycle-dependent manner.^102^ There are 165 ubiquitin ligases encoded in the worm genome;^103^ thus, identifying the specific regulators that control DAF-16 levels in the early embryo may take substantial effort. It is also important to note that, because the cytoplasmic levels of endogenously tagged DAF-16::GFP were nearly equivalent to the background autofluorescence levels innate to worm embryos (*e.g.*, see **Fig. 1B-C**), we cannot rule out DAF- 16 nuclear exclusion in somatic cells. Better tools to continuously track DAF-16 levels in individual embryos will help elucidate the cell and molecular mechanisms that underly DAF-16 patterning in early development.

Why does strong co-disruption of *daf-16* and *daf-2* lead to embryonic lethality? We predict that DAF-2 and DAF-16 both impact cell cycle checkpoint function and/or the mitotic spindle or DNA repair machinery to ensure genome stability in the early *C. elegans* embryo. In support of this model, mitotic progression in germline stem cells is slowed in *daf-2* single mutants, but not in *daf-16; daf-2* double mutants.^104^ Similarly, depletion of the NDC80 complex kinetochore protein, NUF2 (HIM-10 in *C. elegans*), leads to sterility in *daf-2* mutants, but not in *daf-16; daf-2* double mutants.^105^ And inhibition of DNA repair gene expression (by *cdk-12* or *cyclinD/cyd-1* RNAi) promotes germline meiotic arrest in *daf-2* single mutants, but not when *daf-16* and *daf-2* are co-disrupted.^105,106^ These data suggest that DAF-16 may be required for efficient spindle and/or DNA damage checkpoint activation and arrest in the germline. Indeed, ionizing radiation leads to increased oocyte chromatin fragmentation and decreased embryo hatching upon co-disruption of *daf-16* and *daf-2* compared to in *daf-2* single mutants,^107^ as would be predicted if the DNA damage or spindle checkpoint machinery is compromised by loss of DAF-16 activity. Interestingly, in the absence of genotoxic stress, co-disrupting *daf-16* with the milder *daf-2(e1370)* allele,^27^ which has a point mutation in the kinase domain,^85^ does not lead to embryonic lethality.^29^ This suggests that DAF-2 may play a kinase-independent role, like in human preadipocyte cells, wherein insulin receptor/INSR-mediated regulation of mitotic, cell cycle, and DNA damage repair gene expression depends on the intracellular domain but is independent of both insulin binding and kinase activity.^108^

To our knowledge, this is the first report of early cleavage stage embryonic patterning for a FoxO family member in any model system. FoxO genes are expressed at least at the mRNA level in very early cleavage stage *Xenopus tropicalis*,^109^ *Xenopus laevis*,^110^ and Zebrafish embryos.^111^ In female mice, FoxO1/3/4 proteins are expressed in oocytes and early embryos, although they mostly localize to the cytoplasm and perinuclear region until the blastocyst stage, when all three FoxO proteins show nuclear localization in cells of the inner cell mass.^112^ Nevertheless, the injection of siRNAs that target any of the *FoxO1/3/4* genes into 2-cell mouse embryos leads to developmental arrest and prevents blastocyst formation when cultured *in vitro*,^113^ suggesting FoxO family members may be critically important for early murine cleavage divisions. Future studies will be needed to determine if FoxO family members function independent of canonical insulin receptor/INSR signaling in vertebrate embryo development.

## Materials and Methods

Much of these materials and methods are from^65^ with some adaptations. Wormbase (wormbase.org^114^) was used extensively in this work.

### Worm strain maintenance

The *C. elegans* strains used in this study are indicated in **Table S1** and were grown similar to as described^115^. Briefly, worm strains were grown on non-vented standard 60 mm plates (T3308, Tritech Research). Each plate was filled with 10.5 mL nematode growth media (NGM) (23 g Nematode Growth Medium (Legacy Biologicals, a division of Research Products International), 1 mL 1M CaCl_2_, 1 mL of 1M MgSO_4_, 25 mL of 1M K_3_PO_4,_ 975 mL ddH_2_O) using a PourBoy 4 (PB4, Tritech Research) and seeded with 500 μL OP50 *E. coli* bacteria as a food source. Strains were maintained at 20°C, except for JCC2014, which was maintained at 16°C, in cooling/heating incubators (Binder).

### RNA-mediated interference (RNAi) by injection

In general, RNAi-mediated depletion works well in early worm embryos (>95%) because germline protein and mRNA turnover is independent of individual protein and mRNA half-life due to the continual packaging of the syncytial germline into oocytes (*e.g.*, see^116^). For all RNAi experiments, dsRNA was injected directly into L4 stages hermaphrodites 24 hours prior to imaging; uninjected worms transferred at the same time were used as controls. For each target gene, ∼500-1000 bp of gene sequence was PCR amplified using primers containing a T7 promoter sequence and cDNA as a template (when possible, this region was within a single exon). Primers used to generate dsRNA for each target gene are listed in **Table S1**; E-RNAi^117^ and Primer3^118^ were used to design primers. PCR products were confirmed with a 1% agarose gel and PCR purified (QIAquick PCR Purification kit, QIAGEN). The dsRNA was synthesized using a T7 reverse transcription reaction kit (MEGAscript, Life Technologies) and purified using phenol-chloroform. For phenol-chloroform purifications, the synthesized ssRNA was mixed 1:1 with phenol-chloroform (Invitrogen), vortexed for 2 minutes, and spun down for 3 minutes at 12,000 x g in a Beckman Coulter microfuge 16. The aqueous layer was then transferred to new tube, mixed again 1:1 with phenol-chloroform, vortexed for 2 minutes, and spun down for 3 minutes at 12,000 x g in a Beckman Coulter microfuge 16. The aqueous layer was transferred to a new tube, mixed 1:1 with pre-chilled (−20°C) isopropanol (100%, Sigma), and incubated at −20°C overnight (∼16 hours). The ssRNA was precipitated by spinning the tube down at 12,000 x g for 15 minutes in a Beckman Coulter microfuge 16. The pellet (translucent/clear/white) was allowed to air dry for ∼5 minutes and then resuspended in 1x soaking buffer (32.7 mM Na_2_HPO_4_, 16.5 mM KH_2_PO_4_, 6.3 mM NaCl, 14.2 mM NH_4_Cl). To make dsRNA, ssRNA reactions were combined and annealed at 68°C for 10 minutes followed by 37°C for 30 minutes. The dsRNAs were diluted to a final concentration of ∼2000-2500 ng/μL (when possible) and 2 μL aliquots of the dsRNA were stored at −80°C until needed.

For each experiment, a fresh aliquot was diluted to ∼1000 ng/μL using 1x soaking buffer and centrifuged at 16,000 x g for 10 minutes at room temperature (∼22°C) in a Beckman Coulter microfuge 16 prior to use. Machine pulled borosilicate glass capillary needles were made fresh each week (World Precision Instruments capillaries, WPI; Sutter Instruments, P1000 needle puller). 0.4 μL of the diluted dsRNA was loaded into the back of pulled borosilicate glass capillary needles and injected into the gut of L4 hermaphrodites using a Leica DMIRB microscope equipped with Hoffman optics, a Plan L 20x/0.4 CORR PH (Leica), a rotating stage, and the XenoWorks digital microinjector and micromanipulator injection system (Sutter Instruments). Groups of 5-10 worms were injected at a time, rescued by resuspension in M9 buffer (6 g KH_2_PO_4_, 12 g Na_2_HPO_4_, 10 g NaCl, 0.5 mL 1 M MgSO_4_, ddH_2_O to 2 L) on fresh plates seeded with OP50 bacteria, and allowed to recover for ∼24 hours at 20°C prior to imaging.

### Worm and embryo preparation for live cell imaging

Young gravid adult hermaphrodites were dissected on a high-resolution dissecting microscope (Olympus SZX16 with an Olympus SDF PLAPO 1XPF objective) in M9 buffer. Embryos were mounted on a thin (∼1-2x lab tape thickness) 2% agar pad on a glass slide (VWR VistaVision, 3 inches x 1 inch x 1 mm) using a hand-pulled borosilicate glass capillary pipette (World Precision Instruments, WPI) as a mouth pipette. For most experiments, embryos were clustered on the agar pad using an eyelash tool (single eyelash from M. Mauro glued to the end of a hand-pulled borosilicate glass capillary pipette (WPI)). For time lapse imaging of development and genome integrity assays, embryos were clustered on the agar pad using an ultrafine single deer hair with handle (Ted Pella, Inc). To image the germline, oocytes, and intestinal cells in adults (**Fig. S1**), worms were first anesthetized in 1x levamisole (0.01%) for ∼10 minutes prior to mounting on the agar pad. A 22 x 22 mm No. 1.5 glass coverslip (VWR) was placed on top of the embryos or worms (germlines/oocytes/intestine) for imaging, similar to as described^119^.

### Live cell imaging microscope set up and temperature control

For all imaging experiments, room and microscope temperatures were continuously monitored using a Bluetooth-enabled smart temperature sensor (SensorPush) on the microscope stage. All imaging was done at ∼26 ± 0.5°C unless noted. All spinning disc confocal microscope systems were controlled using MetaMorph software (Molecular Devices).

For all experiments (except PIE-1::GFP, intestinal DAF-16::GFP, and most mCh::H2B genome stability analysis), we used an inverted Nikon Eclipse Ti microscope with a spinning disc confocal unit (Yokogawa, CSU-10 with Borealis upgrade (Spectral Applied Research)), a charge-coupled device (CCD) Orca-R2 camera (Hamamatsu Photonics), and a Piezo-driven (Applied Scientific Instrumentation, ASI) motorized stage for Z-sectioning. Focus was established and maintained using Nikon’s “Perfect Focus” before each Z-series was acquired. Two (488 nm and 561 nm) 150 mW Excitation lasers (ILE-2, Spectral Applied Research) were used and controlled by an acousto-optic tunable filter (Spectral Applied Research). A filter wheel (Sutter Instruments) was used for emission filter (525/50 nm and 620/60 nm bandpass (Chroma)) and DIC analyzer selection. Room temperature was established and maintained using a heat pump-based temperature control device (MHWX, MultiAqua).

For the embryonic PIE-1::GFP (**Fig. S3A**), intestinal DAF-16::GFP (**Fig. S4**), and some mCh::H2B genome stability analysis experiments (some of **Fig. 7A-B**), images were acquired on an inverted Nikon Eclipse Ti microscope (modified for compatibility with near-infrared light, as in ^120,121^). The microscope was equipped with a spinning disc confocal unit (Yokogawa, CSU-10 with Borealis upgrade (Spectral Applied Research)), an Orca-R2 CCD camera (Hamamatsu Photonics), and a Piezo-driven (ASI) motorized stage for Z-sectioning. Two solid state (Cairn) 150 mW 488 nm and 561 nm lasers were used for excitation light, with a filter wheel (Ludl Instruments) that was used for DIC polarizer and emission filter (525/50 nm and 620/60 nm bandpass (Chroma)) selection. Room temperature was established and maintained using a heat pump-based temperature control device (Mr. Slim, MSZ-D36NA; Mitsubishi).

### Live cell imaging and analysis parameters

All data analysis was performed using FIJI (FIJI is Just ImageJ) software ^122^.

### Quantitative analysis of average DAF-16::GFP nuclear levels in embryos and oocytes

For quantitative imaging of average nuclear DAF-16 levels, we generated a strain expressing endogenously-tagged DAF-16::GFP^41^ and transgenic mCherry::H2B (mCh::H2B^86^). Embryos were monitored every 60 seconds using mCh::H2B from the 4-cell stage until the 8-cell (**Fig. 2B (top)**, **3B**, **3D**, **4A**, **4C, and 5B**) or the 16-cell (**Fig. 2B (bottom)**) stage, except in **Fig. 1**, in which random groups of embryos at different stages were collected and imaged. At each time point we acquired an image using a 60x Plan Apo 1.40 N.A. oil immersion objective (Nikon) with 2 x 2 binning and 26 x 1.0 µm Z-sectioning of mCh::H2B to determine the cell cycle and developmental stage. Once the desired stage was reached (*e.g.*, after division of the P2 or P3 cell), we then acquired a single time point image stack as described above for both mCh::H2B and DAF- 16::GFP. To image oocytes in **Fig. S1**, we acquired images using a 60x Plan Apo 1.40 N.A. oil immersion objective (Nikon) with 2 x 2 binning and 40 x 1.0 µm Z-sectioning of mCh::H2B and DAF-16::GFP.

For image analysis of nuclear DAF-16::GFP levels throughout different developmental stages (**Fig. 1D, 2C-D, S2B**), in oocytes (**Fig. S1B**), as well as the various RNAi depletions (**Fig. 3C, 3E, 4B**), loss-of-function alleles (**Fig. 4D**), and reproductive ages (**Fig 5C**), the Z-plane containing the most “in focus” image was selected and the nucleus was traced using the circle tool in FIJI. Next, a 50 x 50-pixel box was drawn outside of the embryo to measure the average extracellular background (camera background). This value was then multiplied by the area of the nucleus and subtracted from the total fluorescence intensity in the nucleus. This background subtracted value was then divided by the area of the nucleus to calculate the average nuclear fluorescence intensity. That data was then normalized by dividing each data-point by its replicate control average or pooled control average (except for early embryo DAF-16::GFP analysis in **Fig. 1D**, which was normalized to N2 levels) and plotted as superplots^123^ except for except for early embryo DAF-16::GFP analysis in **Fig. 1D** and **S2B** in which only individual data points are shown. See also schematics in **Fig. 2E, S1C** for a visual depiction of how quantitative image analysis of nuclear DAF-16::GFP levels was performed.

### Quantitative analysis of endogenous DAF-16::GFP nuclear levels in intestine

For quantitative imaging of intestinal/gut DAF-16 nuclear levels, we utilized the above-mentioned strain expressing endogenously tagged DAF-16::GFP and mCh::H2B with and without either RNAi knockdown (**Fig. S4A**) or crossed to different loss-of-function alleles for IIS pathway components (**Fig S4B**). After worms were anesthetized in 1x levamisole (0.01%) for ∼10 minutes, they were mounted (see *Worm and embryo preparation for live cell imaging* above) and images were acquired using a 10x LU Plan Fluor 0.30 N.A. objective (Nikon) with no binning (1 x 1) and 25 x 5.0 µm Z-sectioning of DAF-16::GFP.

For image analysis in **Fig. S4C** intestinal/gut nuclei were scored for presence or absence of DAF-16::GFP. Prior to analysis, all images were set to the same minimum/maximum brightness and contrast parameters. Worms with no visible nuclear DAF-16::GFP, were placed into the group “no nuclear localization in intestinal cells”. Worms with few nuclei or low nuclear signal, were placed into the group “weak nuclear localization in intestinal cells”. Worms with many nuclei or high nuclear signal, were placed into the group “strong nuclear localization in intestinal cells”. See zoomed images in **Fig. S4C** for representative example images of each category.

### Quantitative analysis of fluorescently tagged reporters for RNAi knockdown efficiency

To determine the efficiency of different RNAi knockdowns (*daf-16, daf-18*), we imaged strains expressing DAF-16::GFP^41^ and mNG::DAF-18 (Glow Worms project, Amaka Okafor and Daniel Dickinson, Univ. of Texas-Austin) reporters in embryos with and without the indicated RNAis (**Fig. 1B** and **S5B**). We used a 60x Plan Apo 1.40 N.A. oil immersion objective (Nikon) with 2 x 2 binning and 26 x 1.0 µm Z-sections. For oocyte imaging of DAF-16::GFP in **Fig. S1A**, we used a 60x Plan Apo 1.40 N.A. oil immersion objective (Nikon) with 2 x 2 binning and 40 x 1.0 µm Z-sections. To determine *age-1* RNAi efficiency, we utilized a strain expressing AGE-1::GFP^73^ and imaged the gonad and oocytes using a 60x Plan Apo 1.40 N.A. oil immersion objective (Nikon) with 2 x 2 binning and 25 x 2 µm Z-sections.

Quantification of nuclear DAF-16::GFP levels in *daf-16(RNAi)* embryos was performed as described above (see *Quantitative analysis of average DAF-16::GFP nuclear levels in embryos and oocytes*). To measure gonad levels of AGE-1::GFP (**Fig. S5A**), a single center slice was chosen where a large area of the syncytium was devoid of gonad nuclei, and a 50 x 50-pixel box was drawn to measure the average fluorescence intensity. The same 50 x 50-pixel box was then moved outside the worm to measure the extracellular background. The average extracellular fluorescence intensity was multiplied by 2500 (area of 50 x 50-pixel box) and then subtracted from the fluorescence value measured in the gonad syncytium. To quantify the embryonic levels of mNG::DAF-18, a sum projection was generated in FIJI and an oval surrounding the entire embryo was drawn to measure the total fluorescence intensity. A 20 x 20-pixel box was drawn outside the embryo to calculate extracellular background levels. The average extracellular fluorescence intensity was multiplied by the measured area of the whole embryo and then subtracted from each whole embryo fluorescence intensity value.

### Quantitative analysis of PIE-1::GFP levels

For quantitative imaging of cellular PIE-1 levels in P2, P3, and P4 cells during embryo development we used a strain expressing endogenously-tagged PIE-1::GFP^61^ (**Fig. S3**). At each embryonic stage we captured an image using a 60x Plan Apo 1.40 N.A. oil immersion objective (Nikon) with 2 x 2 binning and 26 x 1.0 µm Z-sectioning. For image analysis, a sum projection of the embryo was generated and the entire P2, P3, or P4 cell was traced using the freehand tool in FIJI. To subtract background, a 20 x 20-pixel box was drawn in the cytoplasm of a non-enriched cell (somatic cell in the embryo anterior, ABa, ABax, or ABaxx). The data was then normalized by dividing by the control average for each replicate. See also schematic in **Fig. S3C** for a visual depiction of how quantitative image analysis of PIE-1::GFP levels in germ precursor cells was performed

### Effect of maternal age on DAF-16 patterning

For reproductive aging experiments (**Fig. 5**), L4 hermaphrodites expressing endogenously tagged DAF-16::GFP and mCh::H2B were mated with males for 24 hours at a 3:1 ratio of males to hermaphrodite. After mating, hermaphrodites were singled out onto fresh plates and moved to a new plate every ∼24-48 hours to ensure continual access to food and to distinguish the mated worms from younger progeny.

For quantitative imaging of nuclear DAF-16::GFP levels, embryos were dissected from worms on day 1 or day 6 of adulthood and were monitored every 60 seconds using mCh::H2B from the 4-cell stage until the 8-cell stage. At each time point we acquired an image using a 60x Plan Apo 1.40 N.A. oil immersion objective (Nikon) with 2 x 2 binning and 26 x 1.0 µm Z-sectioning of mCh::H2B to determine the cell cycle and developmental stage. After cell division of the P2 cell, we then acquired a single time point image stack for both mCh::H2B and DAF-16::GFP and images were analyzed as described above (see *Quantitative analysis of average DAF-16::GFP nuclear levels in embryos and oocytes*).

### Brood size and embryonic lethality quantifications

To quantify brood size (**Fig. 6A**), L4 hermaphrodites of each strain were placed individually onto fresh 35 mm NGM plates seeded with OP50 and incubated at 25-25.5°C. Each hermaphrodite was moved to a fresh plate every 24 hours until egg laying ceased. Approximately 24 hours after adult worms were removed from a plate, the total number of larvae and dead eggs on the plate were counted.

To quantify embryonic lethality (**Fig. 6B**), individual hermaphroditic young adults were moved on their first day of adulthood onto fresh 35 mm NGM plates seeded with OP50 and incubated at 25-25.5°C for 4 hours. After 4 hours, each worm was moved to a fresh plate and incubated for another 24 hours before being removed from the plate. Total numbers of larvae and dead eggs on each plate were counted ∼24 hours after the parental hermaphrodite was removed from each plate. Larvae that died shortly after hatching were counted as dead larvae.

### Analysis of embryonic development

To examine embryonic development (**Fig. 6C-E**), worms were grown at permissive temperature (20°C for control and *daf-16(mgDf50)*; 16°C for *daf-2(m65); daf-16(mgDf50)*) and then moved to fresh plates in their first day of adulthood and incubated at 25-25.5°C for at least 4 hours prior to imaging. Embryos were insulated by a thin ridge of Vaseline around the coverslip to prevent the agar pad from drying out, with gaps left in 2 opposite corners to allow oxygen exchange. Embryos were imaged at 25 ± 0.5°C and DIC images were acquired every 4 minutes for 16 hours with a 20x objective, 1 x 1 binning, and 18 x 2 µm Z-sectioning.

### High resolution analysis of genome stability in early embryogenesis

To analyze genomic stability (**Fig. 7A-B**), we used strains expressing a marker for histone (mCherry::H2B). Worms were grown at permissive temperature (20°C for control and *daf- 16(mgDf50)* single mutants, 16°C for *daf-2(m65)*; *daf-16(mgDf50)* double mutants) and then moved to fresh plates in their first day of adulthood and incubated at ∼25-25.5°C for at least 4 hours prior to imaging. 1-8 cell stage embryos were dissected and imaged at ∼25-26.5°C. Images were acquired every 30 seconds using a 60x 1.4N.A. objective, 2 x 2 binning, and 19 x 1.5 µm Z-sectioning.

### Figure preparation

All figures for this manuscript were all generated using Adobe Illustrator CC (Adobe) and all graphs were created in Prism (Graphpad).

### Statistical analysis

See **Table S1** for a detailed list of all statistical tests and p-values. Briefly, all statistical analysis was performed using Prism 9 (Graphpad) or Excel (Microsoft). All RNAi experiments were repeated 2-4 times (experimental replicates). The number of experimental replicates (N), cells/embryos (n; most figures) or adults (n; **Fig. 6B** and **S4A-C** only) and embryos (n’; **Fig. 6B** only) analyzed is indicated in all Figures. For **Figs. 1D, 2C-D, 3B-C, 6A (left)** and **S2B**, a 1-way ANOVA was used. For **Fig. 6A (right)** a mixed effects ANOVA was used. For **Figs. 6B, 6D, 6E (right),** and **7B**, a Fisher’s exact test was used. For **Figs. 6E (left)** and **S4C** a Chi-squared Contingency test was used. For **all other figures**, a Student’s t-test was performed (unpaired). Error bars in all graphs represent the standard deviation (SD). p-values: n.s.=p>0.05, *=p≤0.05, **=p≤0.01, ***=p≤0.001, and ****=p≤0.0001.

## Acknowledgements

We thank all members of the Canman and Shirasu-Hiza labs for their support, feedback, and advice on this work. We thank Elena Lucchetta for generating strain JCC1092 and Adriana Hernandez, Michelle Schmidt, and Travis Owen for making worm plates and reagents. We thank Eunhee Choi, Rebecca Haeusler, Iva Greenwald, and David Gems for helpful discussions and comments. We thank Oliver Hobert’s lab, Craig Mello’s lab, Alex Hajnal’s lab, the Glow Worms project (Amaka Okafor and Daniel Dickinson), and the CGC (funded by the NIH Office of Research Infrastructure Programs (P40 OD010440)) for worm strains. We thank Hyo Taek Kim for color inspiration (Colors of Star Wars series). This work was funded by: NIH R01GM117407 (JCC), R01GM130764 (JCC), NIH R01AG045842 (MSH), NIH R35GM127049 (MSH), European Research Council CoG ChromoSOMe N°819179 (JD), NSF GRFP DGE-2036197 (JTW), and NIH T32DK007328 (JMP). The authors declare no competing financial interests.

## Author contributions

M.S. Mauro, J.M. Peiser, and J.C. Canman conceived of the project and designed all experiments. M.S. Mauro and S.L. Martin made the dsRNAs for RNAi experiments. Most worm strains were made by M.S. Mauro and J.M. Peiser. M.S. Mauro performed and analyzed all experiments except for the PIE-1::GFP imaging, which was done by S.L. Martin, and analyzed by M.S. Mauro; the maternal age experiment which was done by J.M. Peiser and analyzed by M.S. Mauro; and the *daf-16(mgDf50)* and *daf-2(m65)* experiments and analysis which were done by J.M. Peiser. Intestinal cell imaging was done by M.S. Mauro, J.T. Wiles, and J.M. Peiser. M.S. Mauro, J.M. Peiser, M. Shirasu-Hiza, J. Dumont, and J.C. Canman made significant intellectual contributions and helped write or edit the manuscript. M.S. Mauro, J.M. Peiser, and J.C. Canman made the figures.

