## Supplemental Figures 1-5 and Legends for "Canonical insulin-like receptor DAF-2 signaling-independent patterning and role for FoxO/DAF- 16 in early embryos"

**Figure S1:** ***daf-16(RNAi)* greatly reduces DAF-16 levels in the germline and oocytes. A)** Representative images showing endogenously tagged DAF-16::GFP (green) and mCherry::histone2B (H2B; magenta) localization in the *C. elegans* germline and oocytes with and without *daf-16(RNAi)*; black box behind some images for display; scale bar=10 µm. **B)** Graph plotting the average (avg.) nuclear levels of endogenously tagged DAF-16::GFP in control (grays) and *daf-16(RNAi)* (reds) in immature -5+ oocytes normalized (norm.) to average nuclear levels in controls. Small circles indicate the average levels in individual -5+ oocyte nuclei and large circles indicate replicate averages; shading indicates results from the same replicate (*e.g.*, darkest shades in both control and RNAi conditions same replicate). Error bars=SD; N=number of experimental replicates; n=number of nuclei scored for genotype by color; ****=p-value ≤0.0001 (Student's t-test, unpaired; see **Table S1**). **C)** Schematic depicting analysis shown in **B** to measure nuclear DAF-16::GFP levels in -5+ oocytes (dashed brown circle) and average extracellular background signal (orange box); Mb. partitions=plasma membrane partitions (pea green); chromosomes/nuclei (gray); see also Methods.

Mauro et al Figure S1

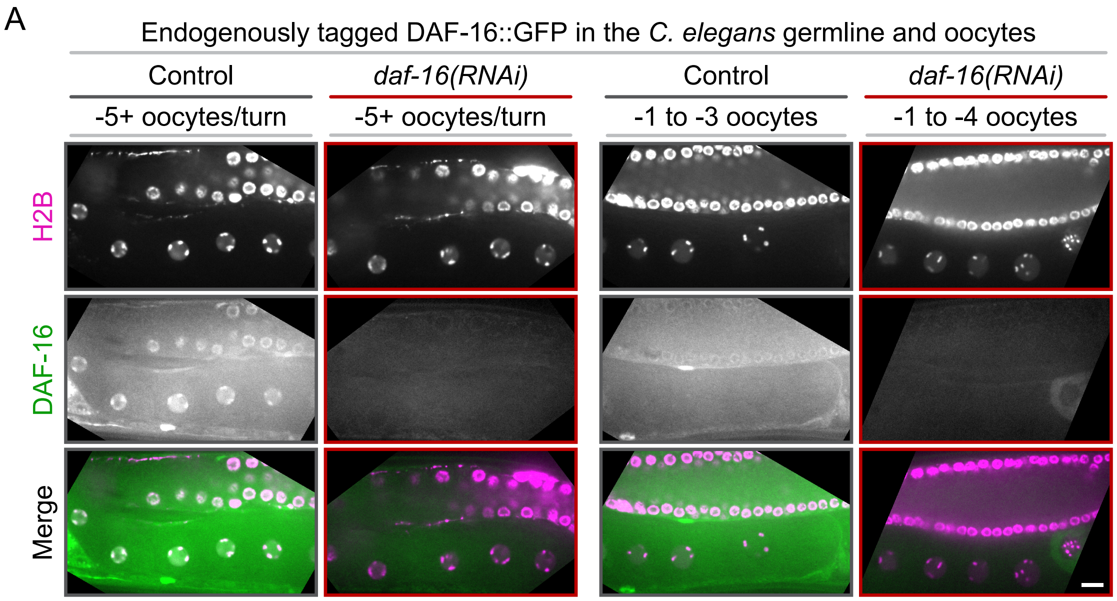


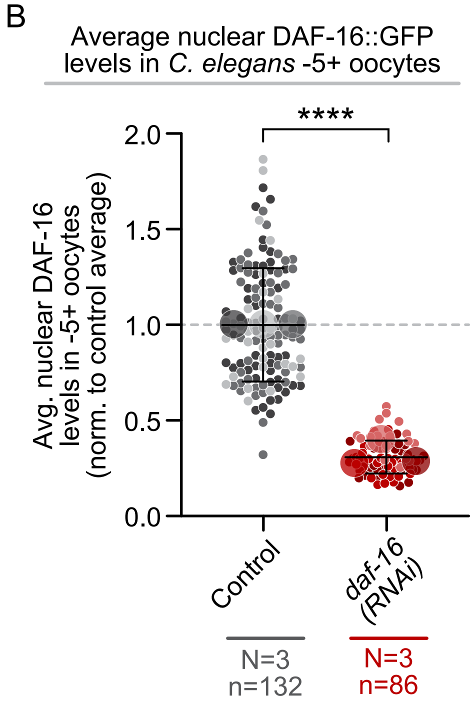


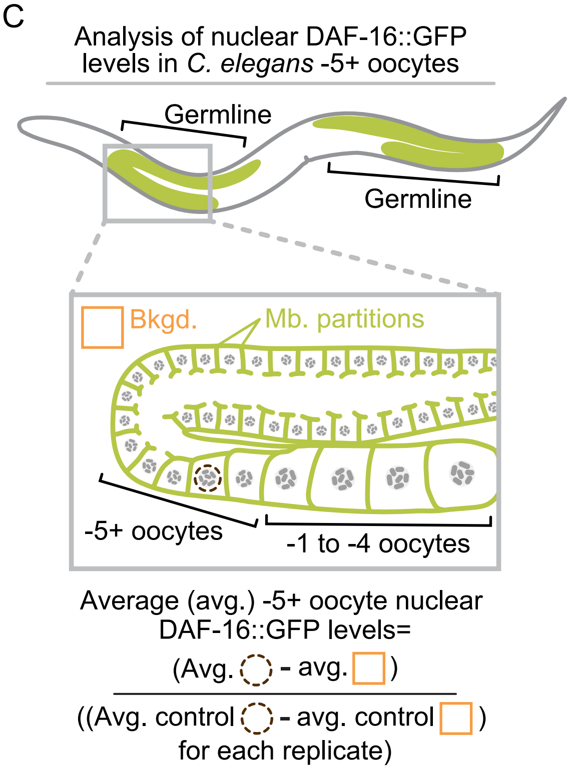

**Figure S2: DAF-16 levels are cell type-independent in 4-cell embryos. A)** Schematic depicting the 4-cell *C. elegans* embryo lineage and each cell identity by color. **B)** Graph plotting average (avg.) nuclear levels of DAF-16::GFP in the ABa (wheat), ABp (olive green), EMS (sage green), and P2 (blue) cells normalized (norm.) to the average (avg.) nuclear levels in the P2 cell. See **Fig. 1B** for representative image. Error bars=SD; N=number of experimental replicates; n=number of nuclei scored for each cell type by color; n.s.=p-value not significant (p-value >0.05) (1-way ANOVA; see **Table S1**).

Mauro et al Figure S2

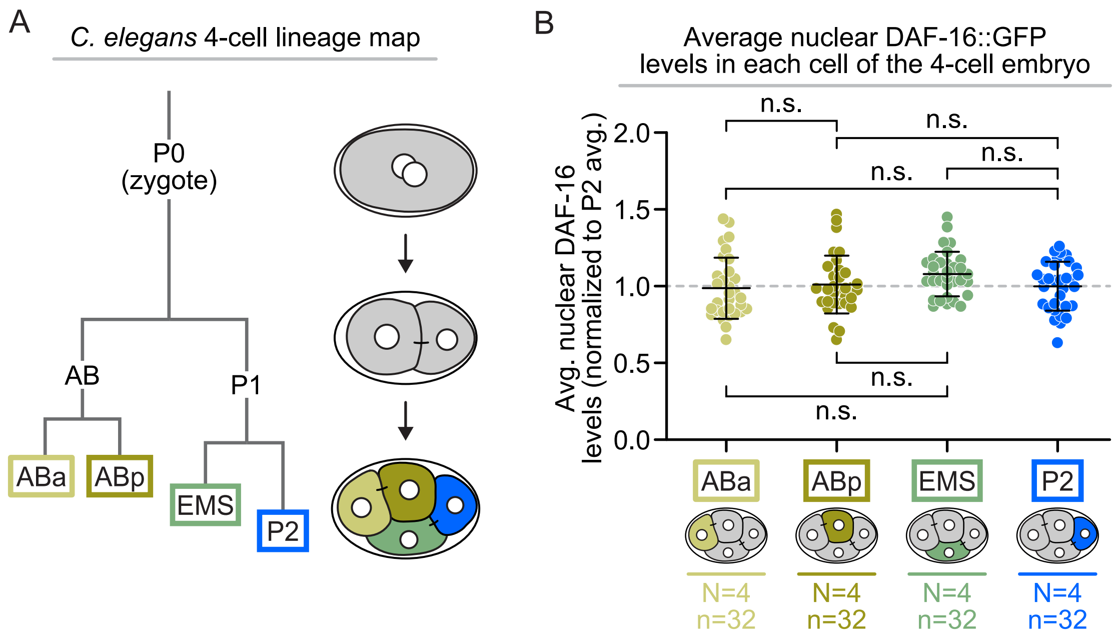

**Figure S3: DAF-16 does not regulate PIE-1 levels in germ precursor cells. A)** Representative maximum projection images showing endogenously tagged PIE-1::GFP localization in the P2, P3, and P4 germ precursor cells (4-cell, 8- to 16-cell, 16- to 32-cell, and 16- to 32-cell over-exposed (to visualize cytoplasmic P granules)) with and without *daf-16(RNAi)*; white dashed line indicates embryo outline; scale bar=10 μm. **B)** Graph plotting the average PIE-1::GFP levels in control (grays) and *daf-16(RNAi)* (reds) in the P2 (left), P3 (center), and P4 (right) germ precursor cells normalized (norm.) to the average (avg.) levels in control (ctrl.) embryos. Small circles indicate the average levels in individual germ precursor cells and large circles indicate replicate averages; shading indicates results from the same replicate (*e.g.*, darkest shades in both control and RNAi conditions from same replicate). Error bars=SD; N=number of experimental replicates; n=number of germ precursor cells scored for each genotype by color; n.s.=p-value not significant (p-value >0.05) (Student's t-test, unpaired; see **Table S1**). **C)** Schematic depicting analysis shown in **B** performed on sum projected images to quantify total PIE-1::GFP levels in individual germ precursor cells and average intracellular background signal in an ABa, ABax, or ABaxx blastomere (orange box); see also Methods.

Mauro et al Figure S3

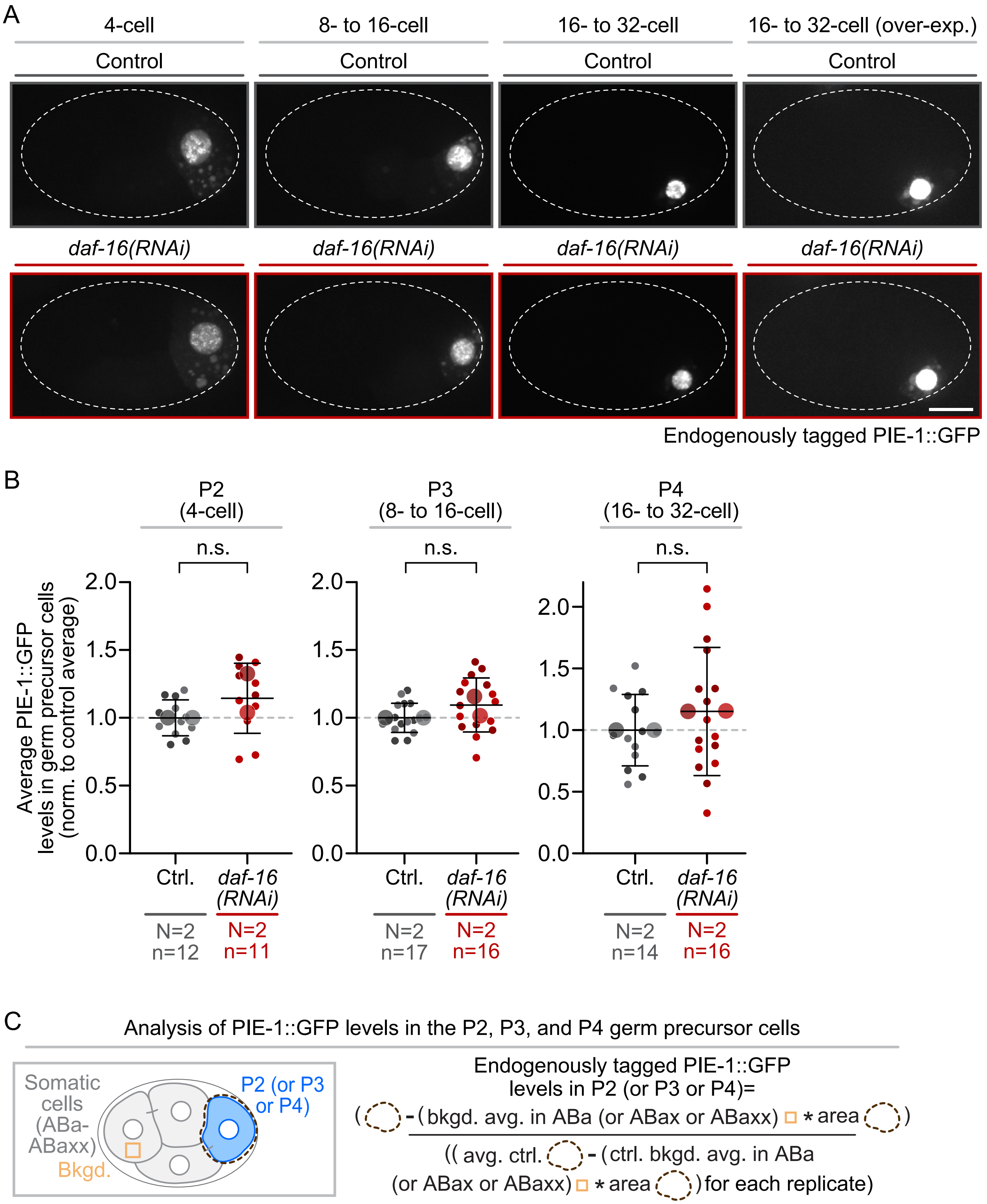

**Figure S4: Intestinal DAF-16 nuclear enrichment with and without RNAi or genetic disruption of IIS pathway components. A)** Representative images of endogenously tagged DAF-16::GFP localization in intestinal cells in control (gray) *daf-2(RNAi)* (green), *age-1(RNAi)* (teal), and *daf-18(RNAi)* (red) adult worms. **B)** Endogenously tagged DAF-16::GFP localization in intestinal cells in control (gray) *daf-2(e1370)* (green), *age-1(hx546)* (teal), and *daf-18(e1375)* (red) adult worms. **C)** Representative images (left) of DAF-16::GFP localization patterns in intestinal cells of adult worms (none=gray, weak=pink, or strong=red) and graphs (right) plotting the percent of worms with each DAF-16::GFP localization pattern after RNAi (top) or genetic disruption (bottom) of IIS components. n=number of worms scored for each genotype by color; *=p-value ≤0.05, ****=p-value ≤0.0001 (Chi-squared Contingency test; only statistically significant comparisons shown on graphs; see **Table S1**). (**A-C**) Orange arrowheads on images indicate intestinal cell nuclei with DAF-16::GFP signal; scale bars=100 μm.

Mauro et al Figure S4

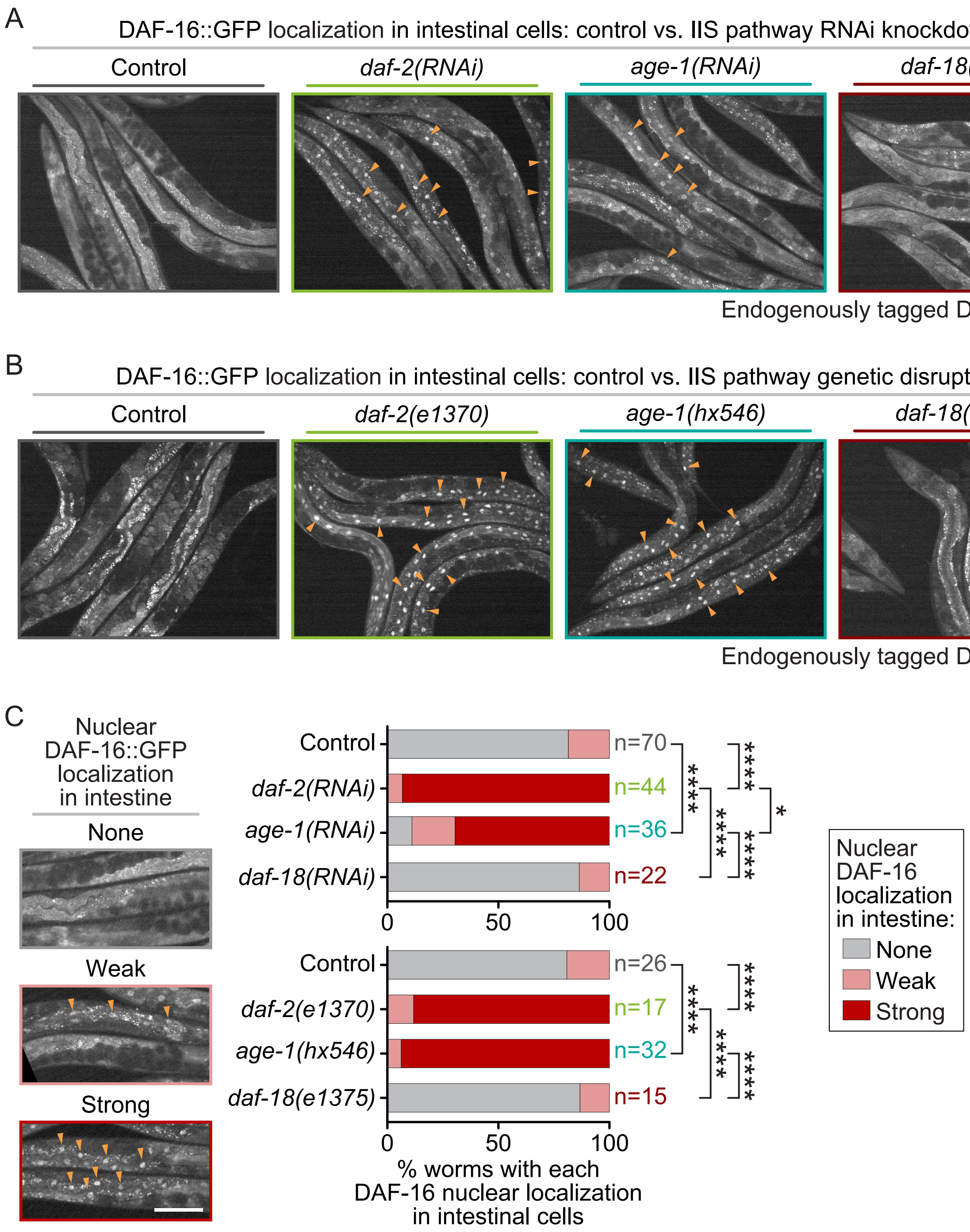

**Figure S5: Depletion of AGE-1 and DAF-18 by RNAi. A)** Representative images (left) of endogenously tagged AGE-1::GFP localization and graph (right) plotting average cytoplasmic levels in adult gonads of control (grays) and *age-1(RNAi)* (teals) worms. **B)** Representative images (left) of endogenously tagged mNeonGreen::DAF-18 (mNG::DAF-18) localization (white dashed line indicates embryo outline) and graph (right) plotting average levels in 8- to 16-cell embryos from control (grays) and *daf-18(RNAi)* (reds) worms. (**A-B**) Black box behind some images for display; scale bars=10 μm. On graphs, small circles=average levels in individual nuclei, large circles=replicate averages, shading=results from same replicate (*e.g.*, darkest shades from same replicate); error bars=SD; N/n=number of experimental replicates and germlines/embryos scored per genotype; ****=p-value ≤0.0001 (Student's t-test, unpaired; see **Table S1**).

Mauro et al Figure S5

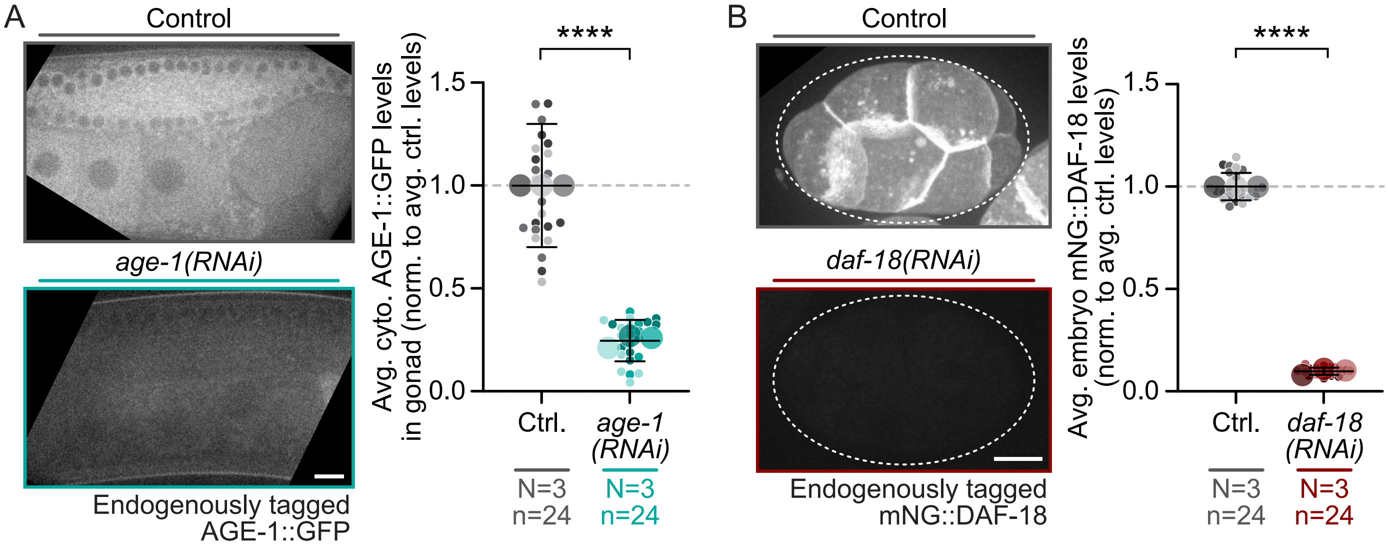
